# Education, intelligence and Alzheimer’s disease: Evidence from a multivariable two-sample Mendelian randomization study

**DOI:** 10.1101/401042

**Authors:** Emma L Anderson, Laura D Howe, Kaitlin H Wade, Yoav Ben-Shlomo, W. David Hill, Ian J Deary, Eleanor C Sanderson, Jie Zheng, Roxanna Korologou-Linden, Evie Stergiakouli, George Davey Smith, Neil M Davies, Gibran Hemani

## Abstract

**Objectives:** To examine whether educational attainment and intelligence have causal effects on risk of Alzheimer’s disease (AD), independently of each other.

**Design:** Two-sample univariable and multivariable Mendelian Randomization (MR) to estimate the causal effects of education on intelligence and vice versa, and the total and independent causal effects of both education and intelligence on risk of AD.

**Participants:** 17,008 AD cases and 37,154 controls from the International Genomics of Alzheimer’s Project (IGAP) consortium

**Main outcome measure:** Odds ratio of AD per standardised deviation increase in years of schooling and intelligence

**Results:** There was strong evidence of a causal, bidirectional relationship between intelligence and educational attainment, with the magnitude of effect being similar in both directions. Similar overall effects were observed for both educational attainment and intelligence on AD risk in the univariable MR analysis; with each SD increase in years of schooling and intelligence, odds of AD were, on average, 37% (95% CI: 23% to 49%) and 35% (95% CI: 25% to 43%) lower, respectively. There was little evidence from the multivariable MR analysis that educational attainment affected AD risk once intelligence was taken into account, but intelligence affected AD risk independently of educational attainment to a similar magnitude observed in the univariate analysis.

**Conclusions:** There is robust evidence for an independent, causal effect of intelligence in lowering AD risk, potentially supporting a role for cognitive training interventions to improve aspects of intelligence. However, given the observed causal effect of educational attainment on intelligence, there may also be support for policies aimed at increasing length of schooling to lower incidence of AD.

## INTRODUCTION

Alzheimer’s disease (AD) is the leading cause of death in England and Wales^1^. Existing treatments are currently unable to reverse or delay progression of the disease. Thus, strategies for reducing the incidence of the disease by intervening on modifiable risk factors are important. Higher educational attainment is associated with a lower risk of dementia^2-5^. However, the mechanisms underlying the associations of educational attainment with AD risk are uncertain and this has implications for intervention design. In particular, what is the role of intelligence? The degree to which education affects intelligence, versus intelligence being largely fixed in early life and acting as a determinant of educational attainment, has been debated for decades^6-10^ and studies have provided evidence of an effect in both directions. ^8 11^ If the principal direction of causality is intelligence to educational attainment, intelligence would induce confounding bias in the association between educational attainment and AD. In this case, interventions aiming to increase educational attainment (e.g. raising the school leaving age to increase years of schooling) are unlikely to affect risk of AD, but alternative prevention strategies such as cognitive training may prove effective. In contrast, if the principal direction of causality is such that greater educational attainment increases intelligence (i.e. intelligence lies on the causal pathway from educational attainment to AD risk), then interventions designed to prolong the duration of education may reduce AD risk, either directly or indirectly through subsequently increasing intelligence.

Determining the relative contributions of education and intelligence to AD risk is of clear importance for designing appropriate policy interventions to reduce AD risk. Using observational methods to unpick these associations is challenging due to bias from measurement error, confounding and reverse causation. More recently, studies have attempted to estimate causal effects of educational attainment on AD risk using methods such as univariable Mendelian randomization (MR). MR is a form of instrumental variable analysis, in which genetic variants are used as proxies for a single environmental exposure^12^. Due to their random allocation at conception, genetic variants associated with a particular risk factor are largely independent of potential confounders, that may otherwise bias the association of interest when using observational methods. Genetic variants also cannot be modified by subsequent disease, thereby eliminating potential bias by reverse causation. Thus, MR can be a useful tool for helping to establish whether the association between an exposure and an outcome is likely to be causal. However, these methods can be problematic with traits that are highly genetically and phenotypically correlated (such as educational attainment and intelligence) ^13 14^. Figure 1 illustrates possible models underlying the observed associations of educational attainment and intelligence with AD risk. In all models shown, causal effects for both exposures on AD risk would be implied from univariable MR analyses. However, depending on the underlying model, intervention targets will differ. Multivariable MR is an extension of univariable MR in which multiple exposures are included within the same model. It can estimate causal effects of one trait, independently of another related trait. Thus, extending MR analyses from the univariable to the multivariable setting may be a useful tool for further disentangling these relationships and establishing the respective roles of both education and intelligence in AD risk ^13^. In this study, we estimated (i) the effect of educational attainment on intelligence and vice versa, (ii) the overall effects of educational attainment and intelligence on risk of AD and (iii) the independent effects of both education and intelligence on risk of AD (i.e. the effects of educational attainment and intelligence on AD risk that are independent of the other trait).

**Figure 1:**
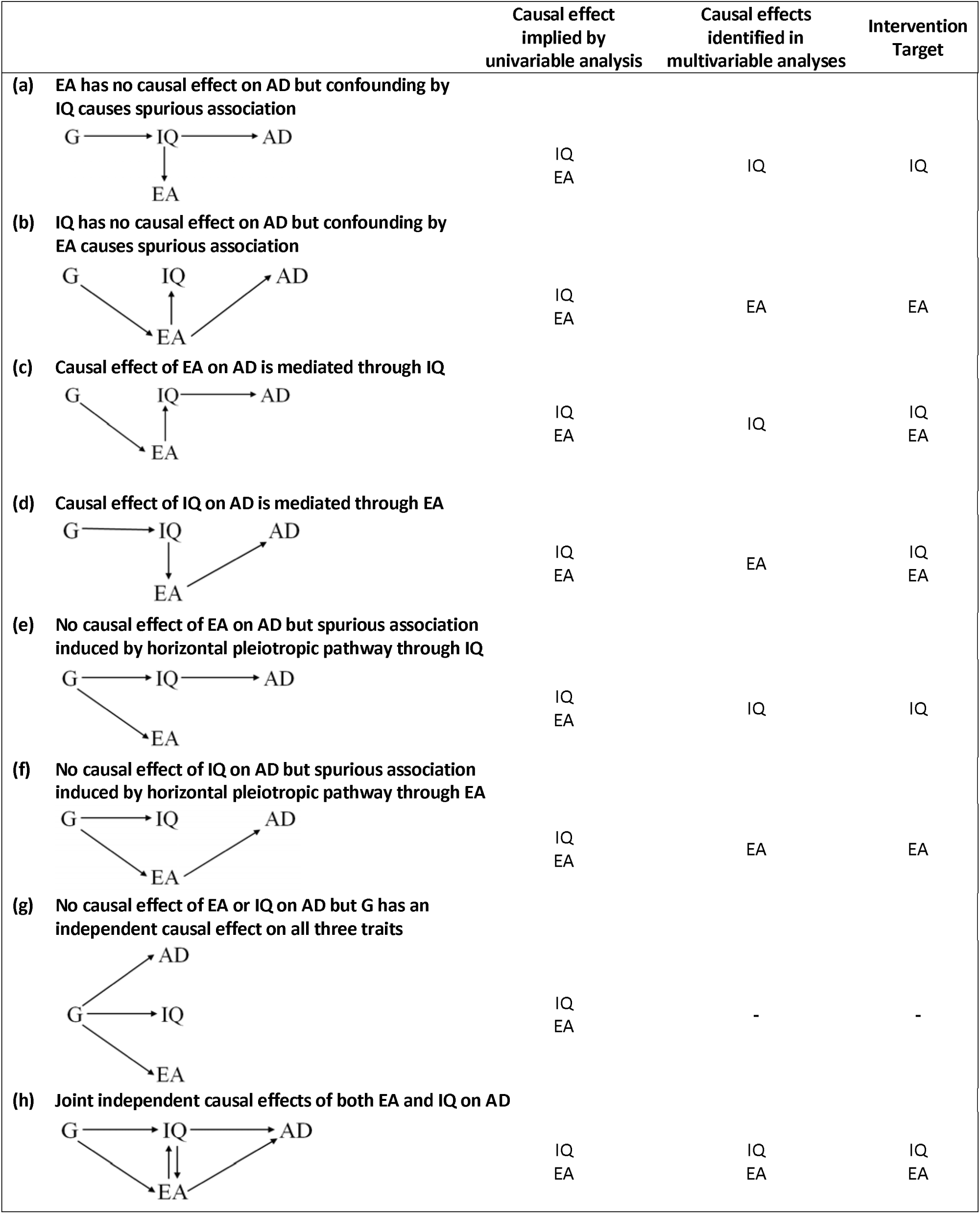
A non-exhaustive list of possible models underlying the observed causal effects of educational attainment, intelligence and risk of Alzheimer’s disease. Please note that these are not intended to be directed acyclic graphs. IQ denotes intelligence. EA denotes educational attainment and AD denotes Alzheimer’s Disease. G denotes a set of instruments, which are drawn as a single node for visual simplicity. Panel (a) illustrates a model in which G is identified in a genome wide association study of EA, because it is associated with EA indirectly through IQ. IQ has an independent effect on AD but EA does not. A spurious association between EA and AD is induced due to confounding by IQ. Accounting for IQ in multivariable analysis would reveal no independent effect of EA on AD risk and the intervention target should be IQ. Panel (b) illustrates a model in which G is identified in a genome wide association study of IQ because it is associated with IQ indirectly through EA. EA has an independent effect on AD but IQ does not. A spurious association between IQ and AD is induced due to confounding by EA. Accounting for EA in multivariable analysis would reveal no independent effect of IQ on AD risk and the intervention target should be EA. Panel (c) illustrates a model in which the effect of EA on AD risk is entirely mediated by IQ (i.e. IQ lies on the causal pathway between EA and AD). Multivariable analyses would reveal an independent effect of IQ on AD risk, but no independent effect of EA. The intervention target could be either IQ or EA. Panel (d) illustrates a model in which the effect of IQ on AD risk is entirely mediated by EA (i.e. EA lies on the causal pathway between IQ and AD). Multivariable analyses would reveal an independent effect of EA on AD risk, but no independent effect of IQ. The intervention target could be either EA or IQ. Panel (e) illustrates a model in which there is full horizontal pleiotropy through IQ. Horizontal pleiotropy occurs when G has a causal effect on disease independently of its effect on the exposure. In this case, multivariate analyses would reveal an independent effect of IQ on AD risk, but no independent effect of EA and the intervention target should be IQ. Panel (f) illustrates a model in which there is full horizontal pleiotropy through EA. Multivariate analyses would reveal an independent effect of EA on AD risk, but no independent effect of IQ and the intervention target should be EA. Panel G illustrates a model in which G independently effects all three traits, but the three traits have no causal effect on each other. Multivariable analysis would show no independent effects of EA or IQ on AD risk. Panel (h) illustrates a model in which there are joint independent effects of both EA and IQ on AD risk. Multivariate analysis would show independent effects of both IQ and EA and the intervention target could be either IQ or EA. Here, the bi-directional relationship between IQ and EA does not affect the qualitative interpretation.

## METHODS

### Mendelian Randomization

MR is a form of instrumental variable analysis that uses genetic variants to proxy for environmental exposures. Two-sample MR^15^ is an extension in which the effects of the genetic instrument on the exposure and on the outcome are obtained from separate genome-wide association studies (GWAS). This method is particularly useful for trying to identify early life risk factors for later life diseases like AD, because unlike in observational studies, rich longitudinal data across the whole life course (which are scarce) are not needed. MR is based on three key assumptions: (i) genetic variants must be robustly associated with the exposure of interest, (ii) genetic variants must not be associated with potential confounders of the association between the exposure and the outcome and (iii) there must be no effects of the genetic variants on the outcome, that do not go via the exposure (i.e. no horizontal pleiotropy) ^16^. To-date, MR studies have typically been univariable (i.e. examining the effect of one exposure on an outcome), thereby estimating the total effect of the exposure on the outcome through all possible pathways. More recently, multivariable MR methods have been proposed to investigate the independent effects of multiple traits on an outcome. Methods for conducting a multivariable MR analysis have been published elsewhere ^13 17 18^

### Data

For educational attainment, we used the GWAS (discovery and replication meta-analysis, n=293,723) ^19^ which identified 162 approximately independent genome-wide significant (p<5×10^−8^) single nucleotide polymorphisms (SNPs) associated with years of schooling. SNP coefficients were per standard deviation (SD) units of years of schooling (SD=3.6 years). For intelligence, we used the largest (n= 248,482) and most recent iteration of the Multi-Trait Analysis of Genome-wide association studies^20^, which identified 194 approximately independent (r^2^ threshold <0.01 within a 10mb window using 1000 genomes reference panel) genome-wide significant SNPs. SNP coefficients were per one SD increase in the intelligence test scores. F statistics provide an indication of instrument strength^22^ and are a function of R^2^ (how much variance in the trait is explained by the set of genetic instruments being used), the number of instruments being used and the sample size. The F statistics for the educational attainment and intelligence instruments are 43.5 and 50.45, respectively (F>10 indicates the analysis is unlikely to suffer from weak instrument bias) ^23^. For the outcome (AD) we used the large-scale GWAS of AD conducted by the International Genomics of Alzheimer’s Project (IGAP, n=17,008 AD cases and 37,154 controls) ^24^. SNP coefficients were log odds ratios of AD. Ethical approval was granted for each of the original GWAS studies and details can be found in the respective publications.

### Estimating the bidirectional association between intelligence and educational attainment

After (i) excluding non-independent SNPs (ii) excluding SNPs that overlapped between the two GWAS and (iii) harmonization across both GWAS, there were 148 genome-wide significant SNPs for educational attainment and 180 for intelligence available for these analyses. Full details of the harmonization procedure are provided in the online supplement. Univariable MR was used to estimate the total effect of intelligence on educational attainment, and educational attainment on intelligence. This was done using inverse-variance-weighted (IVW) regression analysis ^25^. Briefly, IVW regression is where causal effect estimates for each genetic variant are averaged using an inverse-variance weighted formula (taken from the meta-analysis literature) to provide an overall causal estimate of the exposure on the outcome ^26^. In this regression, the intercept is constrained to zero, which makes the assumption of no horizontal pleiotropy. Results are presented in SD units to enable a comparison of the magnitude of effect across both exposures.

### Estimating the total and independent effects of education and intelligence on Alzheimer’s disease

There were 142 genome-wide significant SNPs for educational attainment and 185 for intelligence available for these analyses, after excluding non-independent SNPs and harmonization across both GWAS (full details of harmonization in online supplement). Univariable MR was used to estimate the total effects of both intelligence and educational attainment (separately) on risk of AD, through all possible pathways, using in an inverse-variance-weighted (IVW) regression analysis (described above) ^25^. As mentioned previously, this univariable method has been shown to yield biased effect estimates if the genetic instruments being used are non-specific for the hypothesised exposure ^13 14^. Thus, to demonstrate these effects as they would be observed in a typical univariable analyses, we did not exclude the 9 SNPs that overlapped across education and intelligence GWAS. We then used multivariable MR to estimate the independent effects of both educational attainment and intelligence on risk of AD, by including both exposures within the same model^13^. After clumping the full list of SNPs from both the education and intelligence GWAS (to ensure only independent SNPs are included) and restricting to those SNPs (or proxies) found in the AD GWAS, a total of 231 SNPS were available for the multivariable MR analyses (84 for education and 156 for intelligence, 9 of which overlap between both GWAS).

### Sensitivity analyses

Firstly, in the bidirectional analysis between educational attainment and intelligence, we endeavoured to rule out the possibility that the genetic instruments used to proxy for educational attainment are actually instruments for intelligence and vice versa (i.e. we wanted to test that the hypothesised causal direction was correct for each SNP used). To do this we performed Steiger filtering^27^ for each SNP to examine whether it explains more variance in the exposure than it does in the outcome (which should be true if the hypothesised causal direction from exposure to outcome is correct). We then re-ran analyses excluding those SNPs for which there was evidence that it explained more variance in the outcome than the exposure. Secondly, to check that the SNPs do not exert a direct effect on the outcome apart from through the exposure (which would violate a key MR assumption of no horizontal pleiotropy^12^), we compared results from all univariable (both the bidirectional education on intelligence analyses and the analysis of education and intelligence on AD risk) and multivariable IVW regressions to those obtained with MR-Egger regression. In MR-Egger regression, the intercept is not constrained to zero, thus, the assumption of no horizontal pleiotropy is relaxed. ^16 26 28^ The estimated value of the intercept in MR-Egger regression can be interpreted as an estimate of the average pleiotropic effect across the genetic variants. An intercept term that differs from zero is therefore indicative of horizontal pleiotropy, and the causal effect estimate obtained from an MR-Egger regression is adjusted for the degree of pleiotropy detected^16^. Full details of the MR-Egger regression analyses are provided in the online supplement. Thirdly, we conducted a leave-one-out analysis for the univariable models in which we systematically removed one SNP at a time to assess the influence of potentially pleiotropic SNPs on the causal estimates ^29^. If any single SNP was invalid, there would likely be distortion in the distribution of the causal effects estimates. Fourth, in all univariable analysis, we assessed whether causal estimates from different genetic variants were comparable (i.e. heterogeneity) using Cochran’s Q statistic ^16^. Considerable heterogeneity would imply that the MR assumptions may not be valid for all the variants included in the analysis. Finally, funnel plots were generated to enable the visual assessment of the extent to which pleiotropy is balanced across the set of instruments used in each analysis. Symmetry in these plots provides evidence against directional pleiotropy.

## RESULTS

### Bidirectional effects of intelligence on educational attainment, and their influences on AD risk

Using 180 and 148 genetic instruments for intelligence and educational attainment, respectively (and no overlapping SNPs), we found strong evidence of causal effects both of intelligence on educational attainment, and of educational attainment on intelligence (Table 1). However, the magnitude of the effect was over two-fold greater for educational attainment on intelligence compared with intelligence on educational attainment.

**Table 1:** Bidirectional effect of intelligence on years of schooling.

| Total effects | Causal effect estimates |  |  |
| --- | --- | --- | --- |
| | N SNPs | Standardised $\beta$<br>(95% CI) | P |
| Intelligence on years of schooling | 180 | 0.51 (0.49, 0.54) | 1.77e-95 |
| Years of schooling on intelligence | 148 | 1.04 (0.99, 1.10) | 9.36e-80 |

SNP – single nucleotide polymorphism. $\beta$ – beta coefficient. CI – confidence interval. Results are interpreted per one standard deviation increase years of schooling and intelligence test scores.

The main IVW regression using all SNPs from the educational attainment GWAS showed that, with each SD more years of schooling (i.e. ∼3.6 years), the odds of AD were, on average, 37% lower (95% CI: 23% to 49%). Per one SD higher intelligence test score, the odds of AD were, on average, 35% lower (95% CI: 25% to 43%, Figure 2 and Table C of the online supplement).

**Figure 2:**
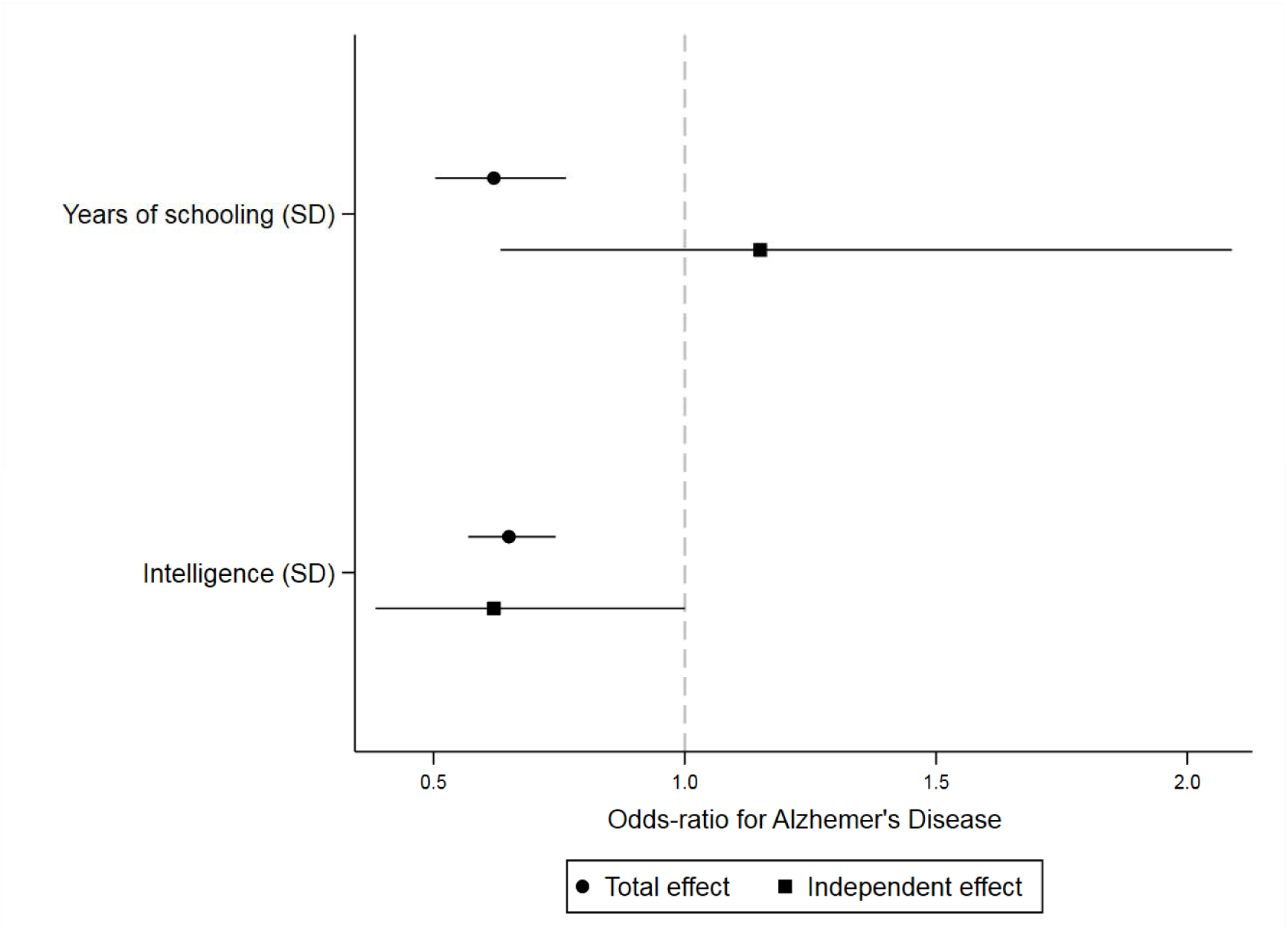
Forest plot showing (i) total effect estimates for years of schooling (in standard deviations) and intelligence (in standard deviations) on odds of AD and (ii) independent effect estimates for both years of schooling and intelligence on odds of AD, when each exposure is adjusted for the other.

### Multivariable analysis of education and intelligence on AD

When both intelligence and educational attainment were included within a single multivariable model, there was little evidence of an effect of educational attainment on AD risk, independent of intelligence (Figure 2 and Table C of the online supplement). There was, however, evidence that higher intelligence lowers risk of AD, independently of educational attainment. On average, after accounting for educational attainment, odds of AD were 38% lower (95% CI: 12% to 56%) per one SD higher intelligence test score Figure 2 and Table C of the online supplement).

### Sensitivity analyses

The Steiger filtering provided evidence that all intelligence SNPs explained more variance in intelligence than educational attainment, suggesting they were all in the correct causal direction (i.e. from intelligence to education). However, there was evidence that 125 (85%) of the 148 education SNPs explained more variance in intelligence than educational attainment, suggesting the hypothesised causal direction is incorrect and is more likely to go from intelligence to education. This left 23 education SNPs. When using only these 23 education SNPs, there was still strong evidence of a causal effect of educational attainment on intelligence (standardised β =0.57, 95% CI: 0.48 to 0.66, Table A of the online supplement), but the magnitude attenuated so that it was comparable to the effect of intelligence on educational attainment (as opposed to the main analysis which showed over 2-fold greater magnitude of effect for education on intelligence than vice versa). There was some evidence of horizontal pleiotropy only in the estimate of the total effect of intelligence on AD risk (Tables B and C of the online supplement). However, for all univariable and multivariable analyses (including the bidirectional effects of intelligence on educational attainment), MR-Egger effect estimates adjusting for pleiotropy were consistently comparable to those from the IWV regressions (Tables B and C of the online supplement). As expected the standard errors were much larger for MR-Egger estimates, because MR-Egger regression provides estimates of two parameters (i.e. both an intercept and a slope) compared to the single parameter in the IVW regressions (i.e. only the slope). The MR-Egger estimate for the total effect of intelligence on risk of AD went in the opposite direction to the IVW estimate (i.e. greater rather than lower odds of AD per SD increase in the intelligence score); however, the confidence intervals were very wide, and the effect estimate could plausibly go in either direction (OR: 1.36, 95% confidence interval: 0.75, 2.48). There was no distortion in the leave-one-out plots for univariable analyses (Figures A to D), suggesting that no single SNP was driving the observed effect from any analysis. There was evidence of heterogeneity in the causal effect estimates from all univariable analyses (P values for all analyses <0.02, Tables B and C of the online supplement). However, provided the pleiotropic effects of genetic variants are equally likely to be positive or negative (i.e. no directional pleiotropy), the overall causal estimate based on all genetic variants is likely to be unbiased and the funnel plots showed little evidence of departure from symmetry (Supplemental figures E to H).

## DISCUSSION

### Bidirectional causal effects in the relationship between of educational attainment and intelligence

In this study we examined the bidirectional effects of intelligence on educational attainment. We found that the relationship between intelligence and educational attainment is indeed likely to be bidirectional in nature (i.e. there is evidence of an effect in both directions), with the magnitude of effect being similar in both directions after filtering SNPs to check they are instrumenting the correct exposure. A recent meta-analysis of quasi-experimental studies of educational effects on intelligence provides evidence that support our MR findings. Across 142 effect sizes from 42 data sets involving over 600,000 participants, the authors reported consistent evidence for beneficial effects of education on cognitive abilities of approximately one to five IQ points (contingent on study design, inclusion of moderators, and publication-bias correction) for an additional year of education ^11^. These findings are similar to ours in respect to magnitude of effect. Assuming a SD of 15 for IQ (as described in the meta-analysis ^11^), intelligence was, on average, up to one-third of a SD higher per year of schooling. In our study we show an average of 0.57 SD higher in intelligence per SD (or. 3.6 years) increase in years of schooling, which equates to 0.16 SD higher intelligence per one additional year of schooling. It is worth nothing that in the quasi-experimental policy reform studies, levels of prior intelligence (or underlying general cognitive ability) will be similar among individuals who left school before and after the policy reforms, making confounding by prior intelligence unlikely. Similarly, in the MR analyses, we endeavoured to exclude any SNPs for education for which there was evidence that they explained more variance in intelligence than education, making it unlikely that our findings for the effect of education on intelligence are a result of all genetic instruments being associated with intelligence and not educational attainment. Thus, both genetic and non-genetic instruments (which contain different sources of bias) provide consistent evidence that educational attainment affects later intelligence. The underlying mechanisms by which educational attainment improves intelligence are uncertain, but several hypotheses have been proposed including the teaching of material directly relevant to the intelligence tests, the training of thinking styles such as abstract reasoning, and the instilling of concentration and self-control^30^. It is also established that learning increases the strength of synaptic connections between neurons in grey matter^31 32^, and human brain imaging has revealed structural changes in white matter after learning complex tasks ^33 34^

Longitudinal observational studies have previously reported associations between early-life intelligence and educational attainment^8^. However, we are unaware of any longitudinal studies that have compared the magnitude of effect for baseline intelligence on educational attainment, with educational attainment on subsequent intelligence in the same sample. One previous study has examined the association between education and lifetime cognitive change after controlling for childhood IQ. The authors reported that (after controlling for childhood IQ score) education was positively associated with IQ at ages 70 and 79 (with the two outcome ages being in different samples), and more strongly for participants with lower initial IQ scores. Education, however, showed no significant association with processing speed, measured at ages 70 and 83 (again, with the two ages being in different samples) ^35^. Another study examined associations between father’s occupation, childhood cognition, educational attainment, own occupation in the 3rd decade, and self-reported literacy and numeracy problems in the 4th decade in the 1946 and 1958 Birth Cohorts^36^. The authors report inverse associations between childhood cognition, educational attainment and adult literacy and numeracy problems. Some studies have looked at genetic overlap between the two traits^20 37^ and reported correlations of up to 0.7^20 38^ but to date, none have explicitly tried to examine the direction of the association using genetic variants that are associated with each of them. As mentioned previously, the largest and most robust evidence to date comes from a recent meta-analysis of quasi-experimental studies of educational effects on intelligence.^11^.

### Effects of educational attainment and intelligence on AD risk

In addition to assessing the bidirectional causal effects in the relationship between educational attainment and intelligence, we also examined the total and independent effects of these traits on risk of AD. Our findings imply that the existing associations reported in the literature between greater educational attainment and lower AD risk are likely to be largely driven by intelligence, rather than there being an independent protective effect of staying in school for longer. This provides evidence against the underlying models illustrated panels (b), (d), (f) and (h) in Figure 1 (i.e. models in which there is an independent effect of educational attainment on AD risk). There are then four main possible explanations for our finding. The first is that prior intelligence is a confounder and induces a spurious association between education and AD risk (i.e. panel (a) in Figure 1). However, given the evidence supporting an effect of education on later intelligence from instrumental variable analyses using policy reforms to increase the school leaving age (in which prior intelligence is randomly distributed among instrument arms and thereby cannot confound), the model in panel (a) is unlikely. The second and third explanations relate to horizontal pleiotropy (either a pathway through IQ as in panel (e) or G independently effecting all traits as in panel (g)). Given our causal effect estimates were comparable when using methods to quantify and adjust for horizontal pleiotropy, these models are also unlikely to fully explain our findings. The fourth explanation is that there is an effect of educational attainment on AD risk, but it is largely mediated by its effects on later intelligence (i.e. panel (c)). Given the existing evidence supporting an effect of education on later intelligence from quasi-experimental studies^11^, and from our own MR analyses, this explanation seems most plausible.

Together, these findings suggest that increasing education attainment (for example, by increasing years of schooling) may have beneficial consequences for future AD incidence. As such, they offer support to the most recent change in school policy in the United Kingdom (in 2013), which now requires young people to remain in at least part-time education until age 18 years (as opposed to 16 years). Our findings also suggest that there may potentially be other ways of reducing risk of AD by improving various aspects of intelligence (e.g. with cognitive training), which may be particularly effective in those with lower educational attainment or in populations where increasing years of schooling is not feasible (e.g. older populations). However, it is worth nothing that it is not clear what type of training (if any) would be beneficial (i.e. memory tasks, abductive reasoning tasks, creative tasks) or when in the life course (and indeed disease course) such training would confer protection (e.g. completing training earlier in life, versus much later but prior to onset of preclinical disease, versus throughout early disease stages).

Our findings are consistent with the ‘brain reserve’ and the ‘cognitive reserve’ hypotheses. Brain reserve refers to structural differences in the brain itself that may increase tolerance of pathology. Cognitive reserve refers to differences in the ability to tolerate and compensate for the effects of brain atrophy, using pre-existing cognitive-processing approaches or compensatory mechanisms ^39^. In support of this, higher levels of education have been shown to be associated with whole brain and ventricular volume as well as cortical thickness ^40-42^. However, it is important to note that these studies often do not consider the potential confounding effects of prior intelligence. One previous study that examined associations between education and brain structure at 73 years found that that the majority of associations observed between education and brain structure (cortical thickness in bilateral temporal, medial-frontal, parietal, sensory and motor cortices) attenuated to the null after accounting for childhood intelligence at age 11, and that neither education nor age 11 IQ was associated with total brain atrophy or tract-averaged fractional anisotropy^43^. A post-mortem study of 130 elderly patients who had undergone cognitive assessment approximately 8 months before death also showed that, at any given level of brain pathology, higher education was associated with better cognitive function^44^. Higher educational attainment may lead to extrinsic compensation through adaptations. Hence, more educated people will usually have occupations that are more intellectually demanding or have greater resources to partake in intellectual activities, resulting in greater cognitive stimulation and consistent with the “use it or lose it” hypothesis ^45^. These compensatory mechanisms may confer protection against advancing AD pathology by increasing the time it takes for an individual to reach the threshold of cognitive impairment, whereby daily living is adversely affected, and a clinical AD diagnosis is made. In addition to compensatory mechanisms, higher education is also associated with avoidance of other potential downstream risk factors such as smoking and excessive alcohol consumption, as well as better engagement with health care systems surrounding primary and secondary prevention (e.g. uptake of and adherence to statin or anti-hypertensive medications).

## Limitations

There are a number of limitations to our study. Firstly, in two-sample MR, “winner’s curse” (i.e. where the effect sizes of variants identified within a single sample are likely to be larger than in the overall population, even if they are truly associated with the exposure) can bias causal estimates towards the null. However, we used SNPs identified in the meta-analysis of the discovery and replication samples of the educational attainment GWAS^19^ making it unlikely that the estimate of the independent effect of education is biased to the null. Secondly, in the presence of weak instruments (i.e. SNPs that are not associated with the exposure at the genome-wide significance level), sample overlap in two-sample MR can bias estimates towards the confounded observational estimate ^46^. There were no overlapping samples in the analysis of educational attainment and intelligence on AD risk, but there was considerable overlap in the samples used for the bidirectional educational attainment on intelligence analysis. Given that all instruments used in the analysis were strong (associated with the exposure at p <5×10^−08^), any bias should be minimal. Thirdly, it is currently not possible to estimate the F statistic (a measure of instrument strength) for multivariable MR in a two- or three-sample setting. Thus, we are unable to assess the conditional strength of our instruments for each exposure, once the SNP effect on the other exposure is taken into account^13^. Fourth, the estimated effect of an exposure on an outcome, that are both associated with mortality, may be susceptible to survival bias^47^. For example, if individuals with lower educational attainment are more likely to die before the age of onset of AD, bias may occur because those individuals with a genetic predisposition for higher educational attainment are likely to live longer, thus having greater risk of being diagnosed with AD. This may induce a non-zero causal effect estimate even if no true biological association exists. In a previous study, we performed simulations to investigate whether our estimates of the effect of educational attainment on AD risk may be biased by survival and found no evidence to suggest this was the case ^5^. Fifth, the phenotype used in the GWAS of intelligence was typically brief (a 2-minute, 13-item test) and heterogeneous. Thus, results may be different if a better phenotype of intelligence was available for GWAS studies. Finally, the educational attainment GWAS only assessed years of full-time academic training from primary education through to advanced qualifications (e.g. degree). Therefore, it remains unclear whether the same genetic variants would be associated with other aspects of education, such as completing vocational courses or completing part-time as opposed to full-time courses. It’s also not clear whether education needs to be completed in a formal setting (such as school or college), or whether any form of learning (e.g. learning new skills ‘on the job’ such as in an apprenticeship during adolescence, or through career development and training courses as an adult in existing full-time employment) would confer the same degree of cognitive protection. This likely depends on the mechanism driving the association between education and AD, thus further studies to unpick the mechanisms may help to shed light on which forms of learning may confer cognitive benefits later in life and in turn, reduce AD risk.

## Conclusions

Our findings imply that there is a bidirectional effect of intelligence on educational attainment and that the magnitude of effect is likely to be similar in both directions. There is robust evidence for an independent, causal effect of intelligence in reducing AD risk. The implications of this are uncertain, but it potentially increases support for a role of cognitive training interventions to improve various aspects of fluid intelligence. However, given that greater educational attainment also increases intelligence, there is potentially also support for policies aimed at increasing length of schooling in order to lower incidence of AD.

## Author contribution statement

ELA, NMD and GH conceptualised the study. ELA completed all statistical analyses with guidance from GH, NMD, LDH and KHW. ELA drafted the first version of the manuscript. All authors provided critical comments on the manuscript. ELA is the guarantor and accepts full responsibility for the work and/or the conduct of the study, had access to the data, and controlled the decision to publish. The corresponding author attests that all listed authors meet authorship criteria and that no others meeting the criteria have been omitted.

## Copyright/Licence for publication

The Corresponding Author has the right to grant on behalf of all authors and does grant on behalf of all authors, a worldwide licence to the Publishers and its licensees in perpetuity, in all forms, formats and media (whether known now or created in the future), to i) publish, reproduce, distribute, display and store the Contribution, ii) translate the Contribution into other languages, create adaptations, reprints, include within collections and create summaries, extracts and/or, abstracts of the Contribution, iii) create any other derivative work(s) based on the Contribution, iv) to exploit all subsidiary rights in the Contribution, v) the inclusion of electronic links from the Contribution to third party material where-ever it may be located; and, vi) licence any third party to do any or all of the above.

